# Trial-By-Trial Auditory Brainstem Response Detection

**DOI:** 10.64898/2026.01.31.703019

**Authors:** George S. Liu, Noor-E-Seher Ali, Dáibhid Ó Maoiléidigh

## Abstract

The neural response of the brainstem to brief sounds, known as the auditory brainstem response (ABR), is widely employed in the laboratory and the clinic to diagnose hearing loss. In contrast to behavioral methods that assess hearing using responses to sounds on a trial-by-trial basis, current ABR approaches are limited to analyzing the *average* ABR over hundreds of trials. Historically, trial-by-trial ABR analysis has not been possible owing to each trial’s small signal-to-noise ratio. Here we overcome this limitation and show how to classify individual ABR trials as detected or undetected. We use the distribution of single-trial ABRs to assess supra-threshold hearing and to define psychophysics-like thresholds, which we call auditory brainstem detection (ABD) thresholds. ABD thresholds decrease as more of the ABR epoch is taken into account, whereas traditional ABR thresholds do not change. Above the ABD thresholds and below 90 dB SPL, signal detection is significantly improved by utilizing more of the ABR epoch. Our method also allows us to rank the supra-threshold hearing ability of individual subjects. Despite having normal ABR thresholds, some subjects appear to have supra-threshold hearing deficits. The trial-by-trial method demonstrates that signal detection by the ensemble of auditory neurons in the brainstem is intrinsically stochastic not only at low stimulus levels, but also at levels up to 100 dB SPL.

**Significance Statement:** Neural responses to sound can be measured by electrodes placed on a subjects head and are commonly used in the laboratory and the clinic to assess hearing. Although the auditory system must distinguish each sound stimulus from intrinsic noise, current methods for analyzing the response of the brainstem to sound only utilize the average response to hundreds of stimuli. Here we overcome this constraint by showing how to classify an individual sound stimulus as detected or undetected based on each auditory brainstem response. This approach can assess hearing at all stimulus levels, indicates that subjects with normal hearing thresholds can exhibit supra-threshold hearing loss, and potentially extends the types of hearing deficits that can be diagnosed using auditory evoked potentials.

## Introduction

The most basic function of hearing is to determine whether a sound is present *each* time a subject is exposed to a sound stimulus. Many behavioral tests address the ability of the auditory system to perform this task (Moore, 2013; Gelfand, 2009). For *each* stimulus presentation interval of a given test, the system classifies a stimulus as being present or absent. Given a fixed false-detection probability, the absolute threshold of hearing can be defined as the sound pressure level at which a certain fraction of stimulus presentations are detected by a subject. In general, a sufficient degree of signal-detection performance for a trial-by-trial classification task defines the absolute threshold of hearing (Macmillan and Creelman, 2005).

Behavioral thresholds are fundamentally different from thresholds based on the auditory brainstem response (ABR). Averaging hundreds of individual ABRs to repeated presentations of the same stimulus reveals a stereotypical temporal pattern (Jewett et al., 1970; Jewett, 1970; Jewett and Williston, 1971; Buchwald and Huang, 1975). The ABR threshold is defined as the sound pressure level at which the *average* ABR evidences the presence of a stimulus (Burkard et al., 2007; Hall, 2007). An extensive array of methods have been developed to objectively determine the ABR threshold (see (Chesnaye, 2019; Suthakar and Liberman, 2019) and references therein). Using the average ABR to define threshold is, however, similar to asking a subject once if they heard a set of stimuli on average *after* all of the stimuli have been presented. In contrast, the absolute threshold of hearing is based on the subject classifying each sound presentation as detected or undetected *before* the next stimulus is presented. Although the ABR threshold is a useful diagnostic, it answers a different question than that addressed by behavioral thresholds.

Supra-threshold hearing can be also assessed using the average ABR. The amplitudes and latencies of characteristic peaks in the response, known as Jewett waves, respectively rise and fall with increasing stimulus levels (Eggermont and Don, 1980; Gorga et al., 1988; Burkard et al., 1990; Kujawa and Liberman, 2009; Lewis et al., 2015). Waves are, however, difficult to identify in all subjects and at low levels (Markand, 1994; Lütkenhöner and Seither-Preisler, 2008). Although the disappearance of a wave may indicate hearing impairment, how to associate changes in wave amplitudes or latencies with supra-threshold hearing loss remains an open question (Kujawa and Liberman, 2009; Lewis et al., 2015; Mehraei et al., 2016; Guest et al., 2018).

Classification of sound stimuli as present or absent is most commonly associated with behavioral thresholds. Here we ask whether trial-by-trial classification of ABRs is possible over the range of physiologically-relevant stimulus levels and propose classification performance as a measure of supra-threshold hearing ability. We also examine whether trial-by-trial classification thresholds, which we call auditory brainstem detection (ABD) thresholds, differ from thresholds based on the average ABR. Our goal is not to find a method that lowers ABR thresholds or that reproduces behavioral thresholds. We seek to evaluate whether the detection question addressed by trial-by-trial classification yields a different answer than that based on the average ABR. Finally, we determine how different ABR quantification procedures change the results based on the average ABR in contrast to trial-by-trial classification.

## Methods

The Administrative Panel on Laboratory Animal Care at Stanford University (APLAC #14345) approved all animal procedures. We measured ABRs in five Sprague Dawley rats at ages p21-p25 using Tucker Davis Technologies (TDT) System 3 RZ6. Rats were anesthetized by administering 3% isoflurane for induction and 1.5-2% isoflurane for maintenance and their body temperatures were held constant at 37*^◦^*C (FHC, DC-temperature controller and heating pad) until they fully recovered. Needle electrodes were placed in the vertex (non-inverting), below the left ear (inverting), and in the ipsilateral hindleg (ground). Subjects were placed in a sound attenuating enclosure within a Faraday cage. Sound stimuli for eliciting ABRs were delivered using a high-frequency speaker (TDT, MF1) that was calibrated using a free-field prepolarized microphone (Model 377C01, PCB Piezotronics) prior to experiments. The rat was positioned 10 cm away from the speaker. The stimulus was a 5 ms sine wave tone pip with cosine-squared envelope rise and fall times of 0.2 ms. The repetition rate was 21 pips per second, the stimulus frequency was 16 kHz, and the intensity was increased in 5 dB steps from 0 dB to 90-100 dB SPL. The sampling rate was 200 kHz, the response was bandpass filtered between 500 Hz and 3 kHz, and artifact rejection was not employed. We recorded responses to 514 stimulus presentations at each sound pressure level for offline analysis. Each response epoch began at the onset of the stimulus, without correcting for the acoustic delay, and had a duration of 7.755 ms. Data is available upon request.

Data analysis was performed in MATLAB (version R2018b, MathWorks). Error bars represent standard errors and, apart from the average features, were estimated by studentized bootstrapping for each stimulus level and rat using 500 resamples for each observed set of feature values. Doubling the number of bootstrapped samples changed the error estimates by less than 10%. P-values, thresholds, cutoff values, false-positive rates, true-positive rates, and standard errors at points that were not directly sampled were estimated by linear interpolation between sampled points.

## Results

### The Average ABR

The average ABR to a stimulus of magnitude *s* presented *N* times is given by

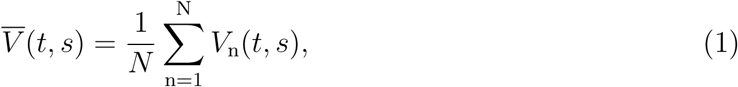

in which *V*_n_(*t, s*) is the response to the *n*^th^ presentation and 0 ≤ *t* ≤ *T*. Each individual ABR *V*_n_(*t, s*) is measured from the onset of the *n*^th^ presentation at *t* = 0 to a time *t* = *T*. Classically, the ABR threshold is the lowest sound pressure level at which the average ABR indicates the presence of the stimulus.

Comparisons between the temporal structures of average ABRs across stimulus levels are facilitated by normalizing *V̄*(*t, s*) by its maximum over time

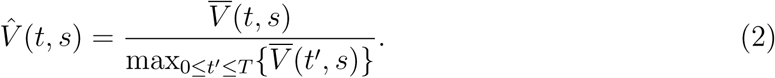

We demonstrate our approach by using the responses of a representative rat to a tone burst. Waves I–IV in this rat are clear in the normalized average ABRs at high stimulus levels, but become more difficult to discern at low levels (Fig. 1). To objectively compare responses at different levels, we choose a summary feature.

**Figure 1:**
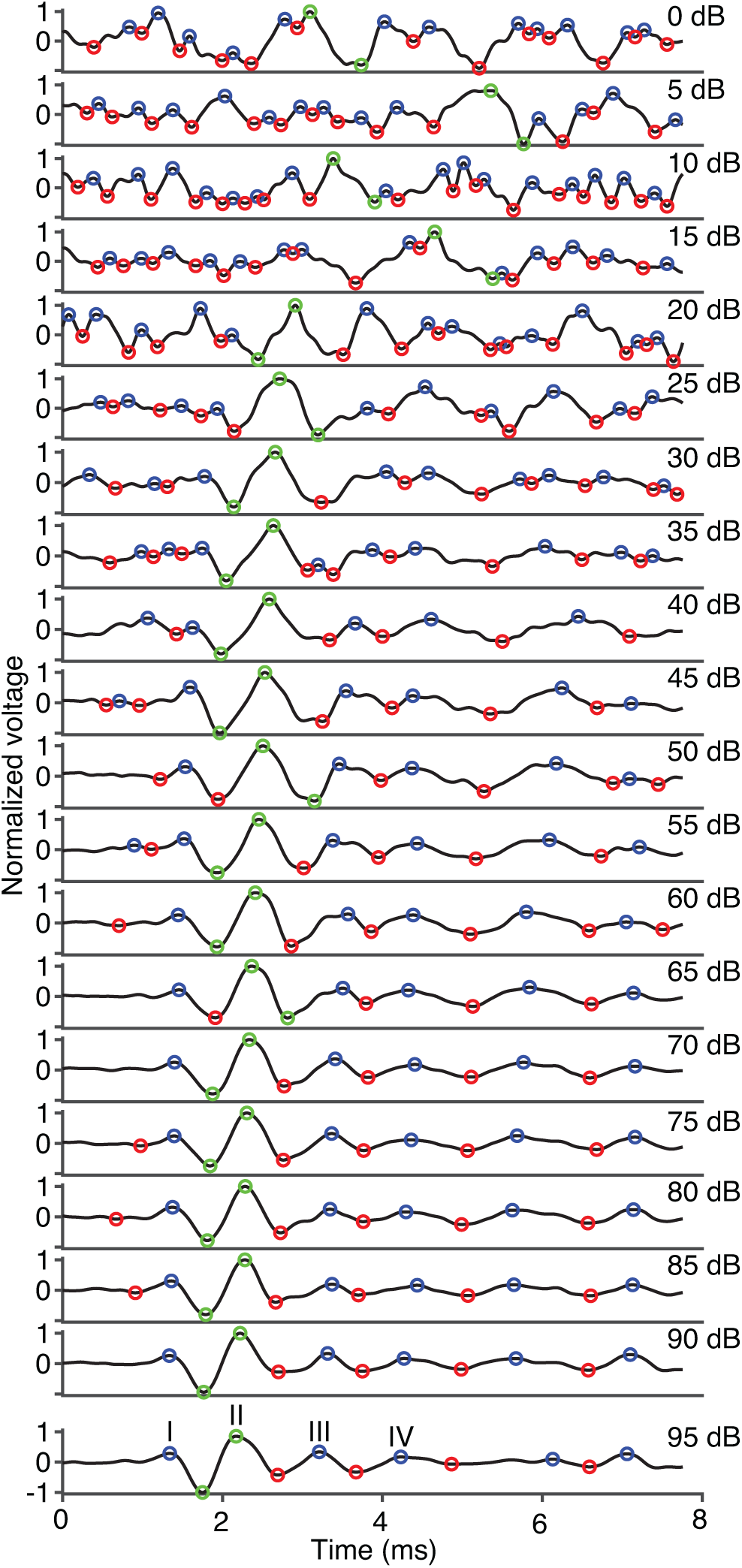
Normalized ABRs of a representative rat are shown at different stimulus levels. We find the peaks (blue circles) and troughs (red circles) of the average ABRs by applying a peak-detection algorithm to the normalized responses with a detection limit of 0.1. The peak and trough corresponding to the largest peak-to-trough difference at each level are indicated by green circles. Jewett waves I-IV are labeled at 95 dB SPL.

### The Voltage-difference Feature

A simple feature that captures the magnitude of the average ABR is the maximum amplitude, measured as the maximum peak-to-trough difference, over the response time *T*. We use a peak-detection algorithm to automatically find the peaks and troughs of the normalized ABR (Jacobson, 2001), allowing us to identify the maximum peak-to-trough difference of the average ABR at each stimulus level

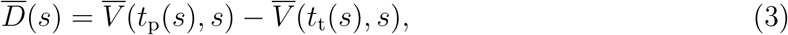

in which *t*_p_(*s*) and *t*_t_(*s*) are respectively the times at which the peak and trough occur for the stimulus level *s*. In the representative rat, *D̄*(*s*) is equal to the difference between wave II and its prior trough at high levels but not at low levels. The feature *D̄*(*s*) is the average of the voltage differences

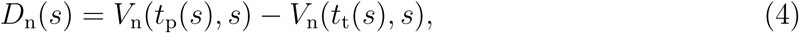

in which *n* ∈ {1*,…, N* }. The distribution of the voltage difference *D* moves to higher voltages as the stimulus level increases, but is not normal at any stimulus level (Q-Q plots indicate heavier tails than normal distributions, Anderson-Darling tests *p <* 5 × 10*^−^*^4^, kurtosis = 5.5 — 14) (Fig. 2).

**Figure 2:**
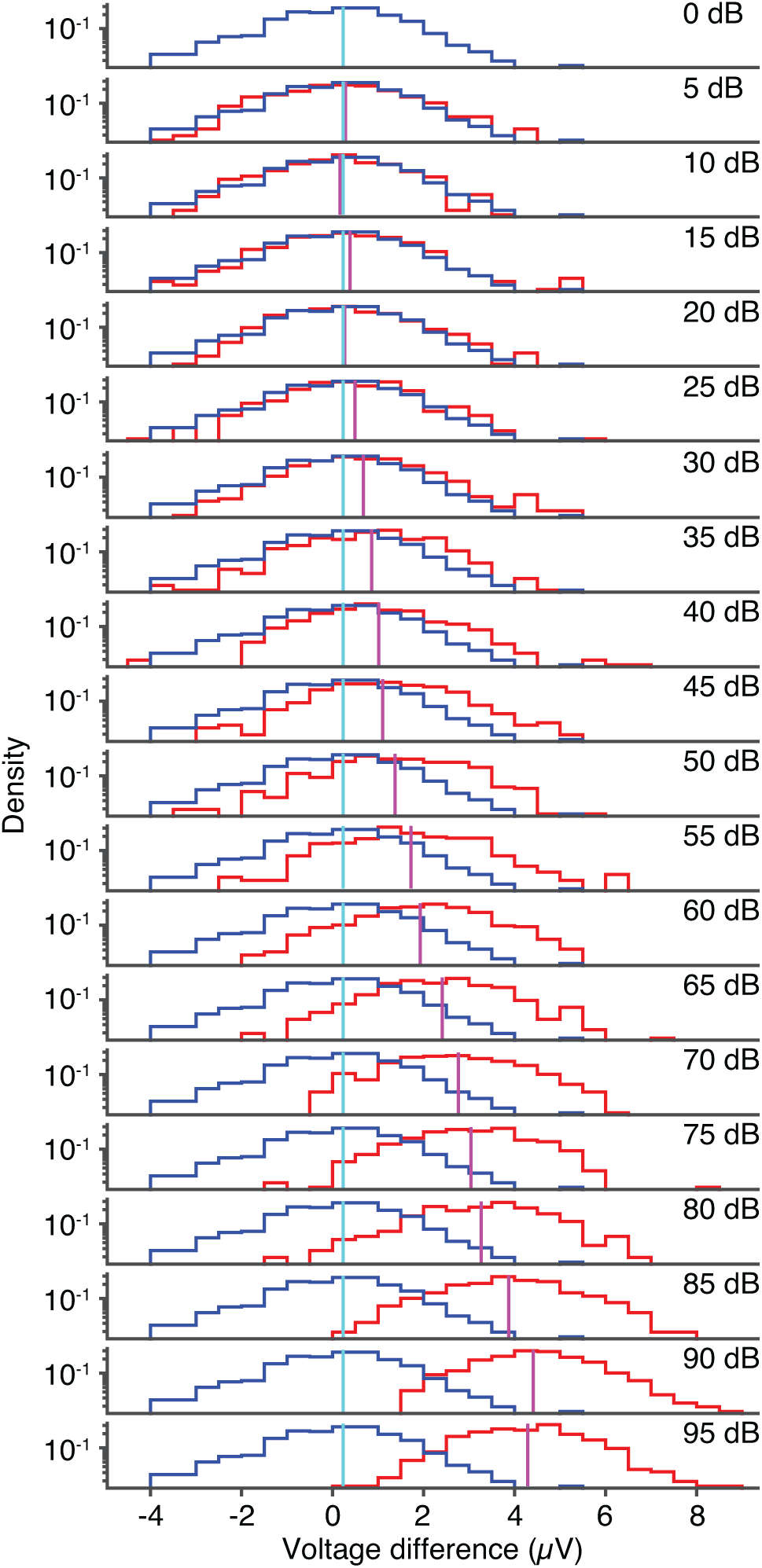
The voltage-difference distribution of the representative rat is shown at different sound pressure levels (red). The distribution’s mean is indicated by a vertical magenta line. The distribution at 0 dB SPL approximates the no-stimulus distribution (blue) and has a mean close to 0 *µ*V (vertical cyan line).

### The ABR Threshold

For a given feature *F*, an ABR threshold can be defined as the stimulus level at which the feature’s distribution owing to stimulation can be distinguished from the distribution in the absence of stimulation using a hypothesis test, in which the null hypothesis is that the distributions are the same. We choose the threshold to be the stimulus level *S* at which a test’s p-value equals 0.05 and for which the p-values at all higher stimulus levels are less than 0.05, that is,

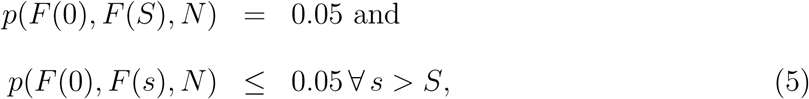

in which the p-value *p*(*F* (0)*, F* (*s*)*, N*) depends on the stimulus level *s* and the number of presentations *N*.

To determine the ABR threshold, we compare *D̄* in the absence of a stimulus to averages arising from the presence of stimuli using Welch’s t-test, in which the *D* distribution in the absence of a stimulus is approximated by the distribution at 0 dB SPL. We use Welch’s t-test which assumes the distributions of the means to be normal without requiring the feature variances to be the same. We note the variance of a feature may change with the stimulus level and the distributions of the means are likely normal owing to the large sample size even though the *D* distributions are not normal (Fig. 2) (Lumley et al., 2002). The ABR threshold defined using this detection rule is 25 dB SPL for the representative rat (Fig. 3(a)). Owing to the standard error of the mean decreasing as the number of stimulus epochs *N* increases, the threshold also declines as *N* rises.

**Figure 3:**
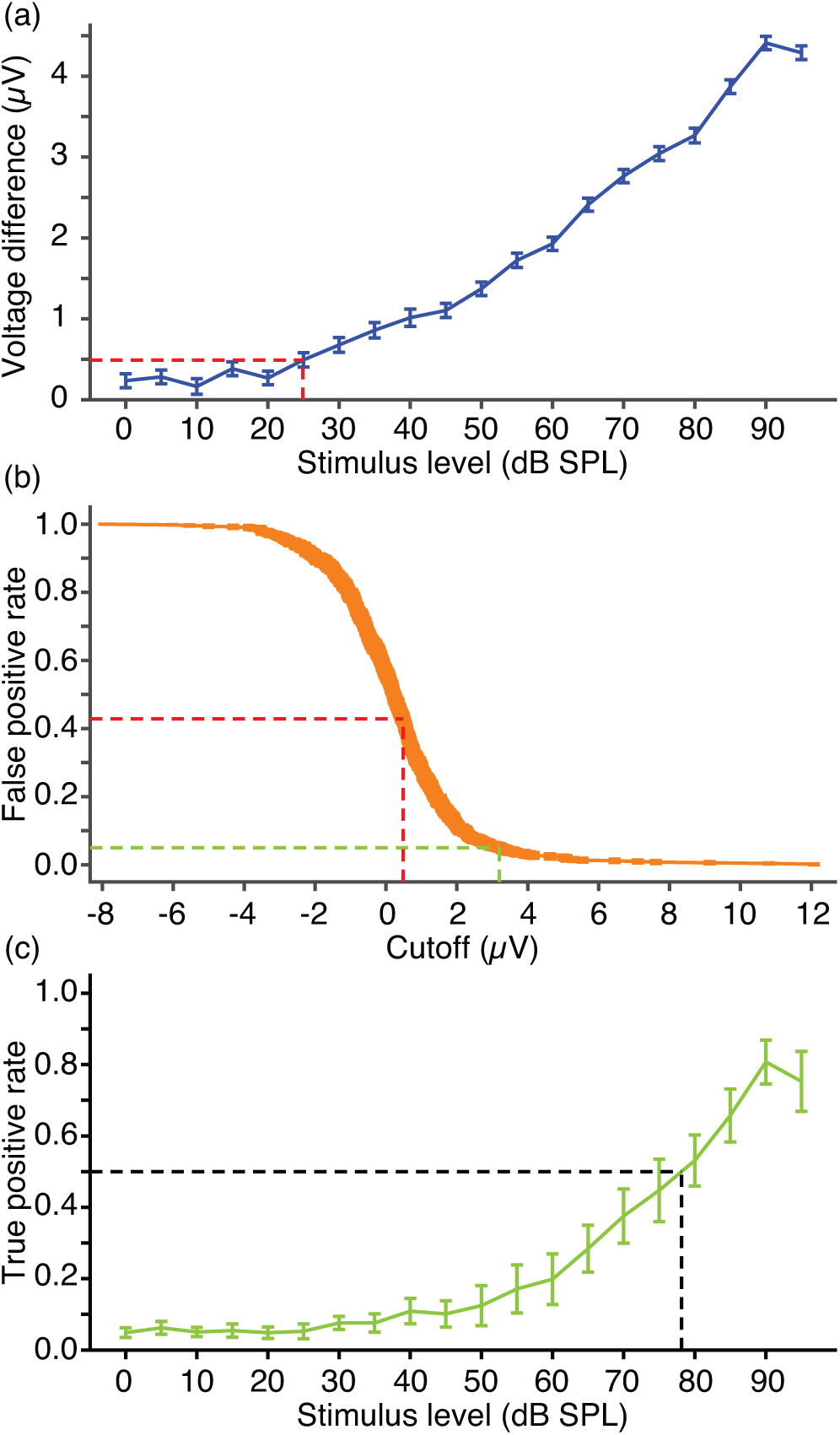
(a) The average ABR’s voltage-difference is shown as a function of the stimulus level and is compared at each level with the 0 dB SPL average using Welch’s t-test. A p-value of 0.05 corresponds to a stimulus level of 25 dB SPL and a cutoff of 0.49 ± 0.09 *µ*V (dashed red lines). (b) The FPR is shown as a function of the voltage-difference cutoff. The cutoff indicated in panel (a) corresponds to a FPR of 0.43 ± 0.02 (dashed red lines). A larger cutoff of 3.2 ± 0.4 *µ*V is needed to achieve a FPR of 0.05 (dashed green lines). (c) The TPR is shown as a function of the stimulus level, in which the voltage-difference cutoff is set to ensure a FPR of 0.05. A TPR of 0.5 determines the PPT of 78 ± 5 dB SPL (black dashed lines).

### The Auditory Brainstem Detection Threshold

Because Eq. 5 uses *all N* stimulus presentations to make a single determination about whether the group of stimuli are detected, it defines a threshold that is fundamentally different in character to the behavioral threshold. The behavioral threshold is found by assessing whether *each* stimulus presentation is perceived (Macmillan and Creelman, 2005; Moore, 2013; Gelfand, 2009). Although an estimate of the behavioral threshold becomes more accurate as the presentation number increases, the behavioral threshold does not depend on this number.

To assess an ABD threshold in a manner analogous to the behavioral threshold, we identify a feature’s distribution with a subject’s internal decision-space distribution (Macmillan and Creelman, 2005). An ABD threshold corresponds to the stimulus level at which there is a sufficiently large difference between a feature’s distribution in the presence and absence of a signal. In contrast to the ABR threshold, which is determined by a statistically-significant difference between averages, an ABD threshold is defined by an effect size. Because a statistically-significant difference is required before determining an effect size, the ABD threshold is expected to be larger than the ABR threshold.

In the case of Welch’s ABR threshold, the effect size for a large number of trials can be estimated by

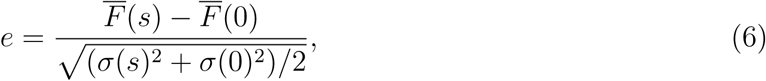

in which *F̄*(*s*) and *σ*(*s*) are respectively the summary feature’s average and standard deviation at stimulus level *s* (Aoki, 2020). When the standard deviation is the same at each stimulus level, the effect size *e* is known as Hedges *g*, Cohen’s *d*, or *d^r^* and is commonly used in psychophysics to define signal-detection performance (Green, 1960; Green and Swets, 1966; Aoki, 2020; Hedges, 1981; Cohen, 1988; Macmillan and Creelman, 2005).

Because the definition of *e* is based on the assumption of normal feature distributions, its interpretation is unclear for unclassified non-Gaussian distributions, such as the ABR distributions discussed here. There are alternative choices for defining an ABD threshold, however, that do not depend on the feature distributions being normal.

### The Positive-pair Threshold

An ABD threshold may be defined by treating the brainstem as a classifier that mimics a subject’s decision process by specifying a cutoff value for the feature, above which an individual ABR is classified as detected. The cutoff determines the false positive rate (FPR)—the fraction of false positive classifications in the absence of a stimulus. For a fixed FPR, the positive-pair threshold (PPT) is defined as the sound pressure level at which the fraction of true positive classifications, the true positive rate (TPR), is sufficiently large. The PPT is one possible choice for defining an ABD threshold. Given a set of decision-space distributions, each positive pair (FPR, TPR) corresponds to a particular sensory sensitivity or discriminability. An acceptable FPR is typically 0.05 or less and the TPR at threshold is usually defined to be 0.5 (Moore, 2013; Macmillan and Creelman, 2005).

The FPR derived from the no-stimulus distribution *P* (*F,* 0) of a feature *F* is given by

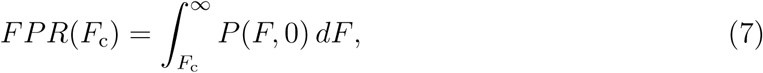

in which *F*_c_ is the feature’s cutoff. For the voltage difference, the FPR decreases as the cutoff *D*_c_ increases (Figs. 2 and 3(b)). The FPR is very high (0.43 ± 0.02) at the cutoff corresponding to the ABR threshold defined using Eq. 5. We can, however, find the feature cutoff such that the FPR is at an acceptable level. Enforcing a FPR of 0.05 defines the cutoff *F*_c5_ according to

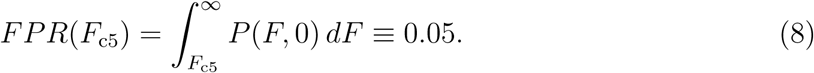

Using this cutoff, the *n*^th^ stimulus presentation at stimulus level *s* is detected if

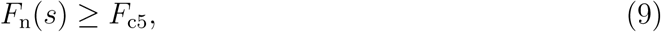

in which *F*_n_(*s*) is the value of the feature for the *n*^th^ stimulus. The TPR is the fraction of times the detection rule Eq. 9 is satisfied and is given by

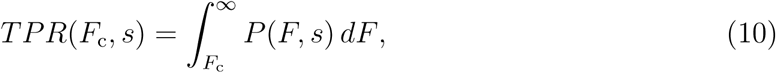

in which *s* /= 0. The TPR grows as the stimulus level rises and we define the PPT to be the stimulus level at which the TPR equals 0.5. The PPT for the representative rat’s voltage difference is 78 ± 5 dB SPL and is based on a choice of feature and the pair (FPR, TPR) (Fig. 3(c)).

### The Inner-product Feature

The ABD threshold can be found for many different types of features, including features whose average cannot be calculated from the average ABR alone. To illustrate, we employ the inner-product feature given by

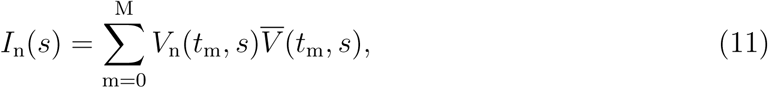

in which *t*_m_ = (*m/M*)*T* are the times at which the ABR is sampled (Coppola et al., 1978). The inner product quantifies the similarity between each response and a template, given by the average ABR, and is similar to the cross-correlation between two deterministic time series. As the stimulus intensity grows, the inner-product distribution widens and shifts to higher values of the inner product, and is non-Gaussian at all stimulus levels (Q-Q plots indicate heavier tails than normal distributions, Anderson-Darling tests *p <* 5×10*^−^*^4^, kurtosis = 7.0 — 12) (Fig. 4).

**Figure 4:**
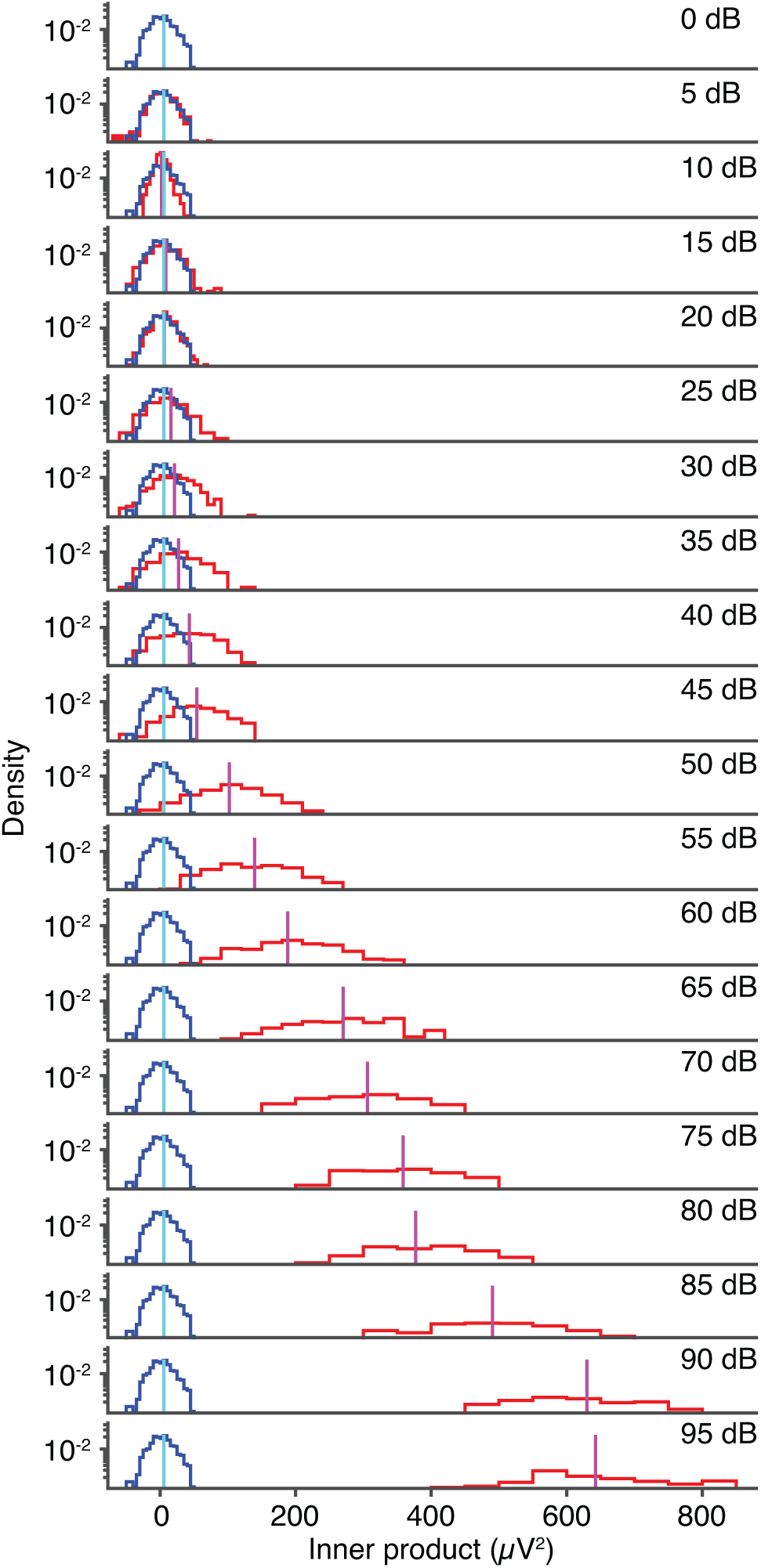
The inner-product distribution of the representative rat is shown at different sound pressure levels (red). The distribution’s mean is indicated by a vertical magenta line. The distribution at 0 dB SPL approximates the no-stimulus distribution (blue) and has a mean close to 0 *µ*V^2^ (vertical cyan line).

As before, we use Welch’s t-test to find the ABR threshold at which the average *Ī*(*s*) at stimulus level *s* can be distinguished from the average in the absence of a stimulus *Ī*(0) (Eq. 5, Fig. 5(a)). The ABR threshold using the inner product is 25 dB SPL, but the FPR for detecting individual stimulus presentations is again high (0.27 ± 0.02) (Fig. 5(b)). Raising the inner product cutoff to ensure a FPR of 0.05 results in a PPT of 40 ± 2 dB SPL (Fig. 5(c)).

**Figure 5:**
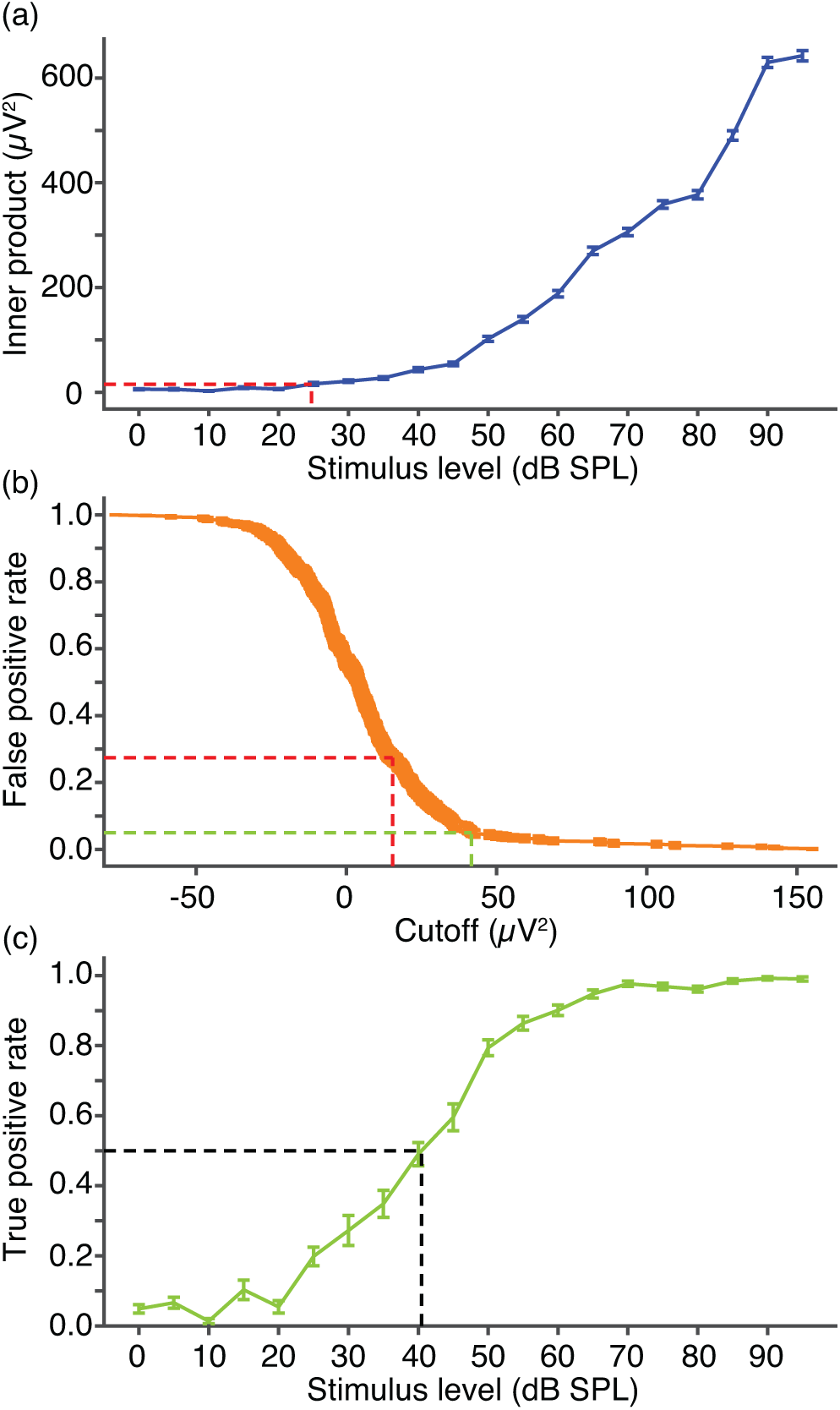
(a) The average inner product is shown as a function of the stimulus level and is compared at each stimulus level with the 0 dB SPL average using Welch’s t-test. A p-value of 0.05 corresponds to a stimulus level of 25 dB SPL and a cutoff of 15 *µ*V^2^ (dashed red lines). (b) The FPR is shown as a function of the inner-product cutoff. The cutoff indicated in panel (a) corresponds to a FPR of 0.27±0.02 (dashed red lines). A larger cutoff of 42±0.4 *µ*V^2^ is needed to achieve a FPR of 0.05 (dashed green lines). (c) The TPR is shown as a function of the stimulus level, in which the inner-product cutoff is set to ensure a FPR of 0.05. A TPR of 0.5 determines the PPT of 40 ± 2 dB SPL (black dashed lines).

### Supra-threshold Signal-detection Performance Depends on the Choice of Feature

Although the ABR thresholds based on the voltage difference and the inner product are similar, the PPTs differ by 40 dB. Whether different features yield the same thresholds evidently depends on the cutoffs employed. To determine how a feature’s performance changes with the cutoff, we find the FPR and the TPR over the range of cutoffs and sound pressure levels (Fig. 6(a, b)). The resulting curves are known as receiver operator characteristic (ROC) curves and they illustrate the ability of the auditory system to detect a stimulus of size *s* if the only information utilized were the value of the feature in question (Green, 1960; Fawcett, 2006). The line of no discrimination is the diagonal of unit slope given by *TPR*(*F*_c_, 0) = *FPR*(*F*_c_). Perfect stimulus detection corresponds to a FPR of zero and a TPR of unity.

**Figure 6:**
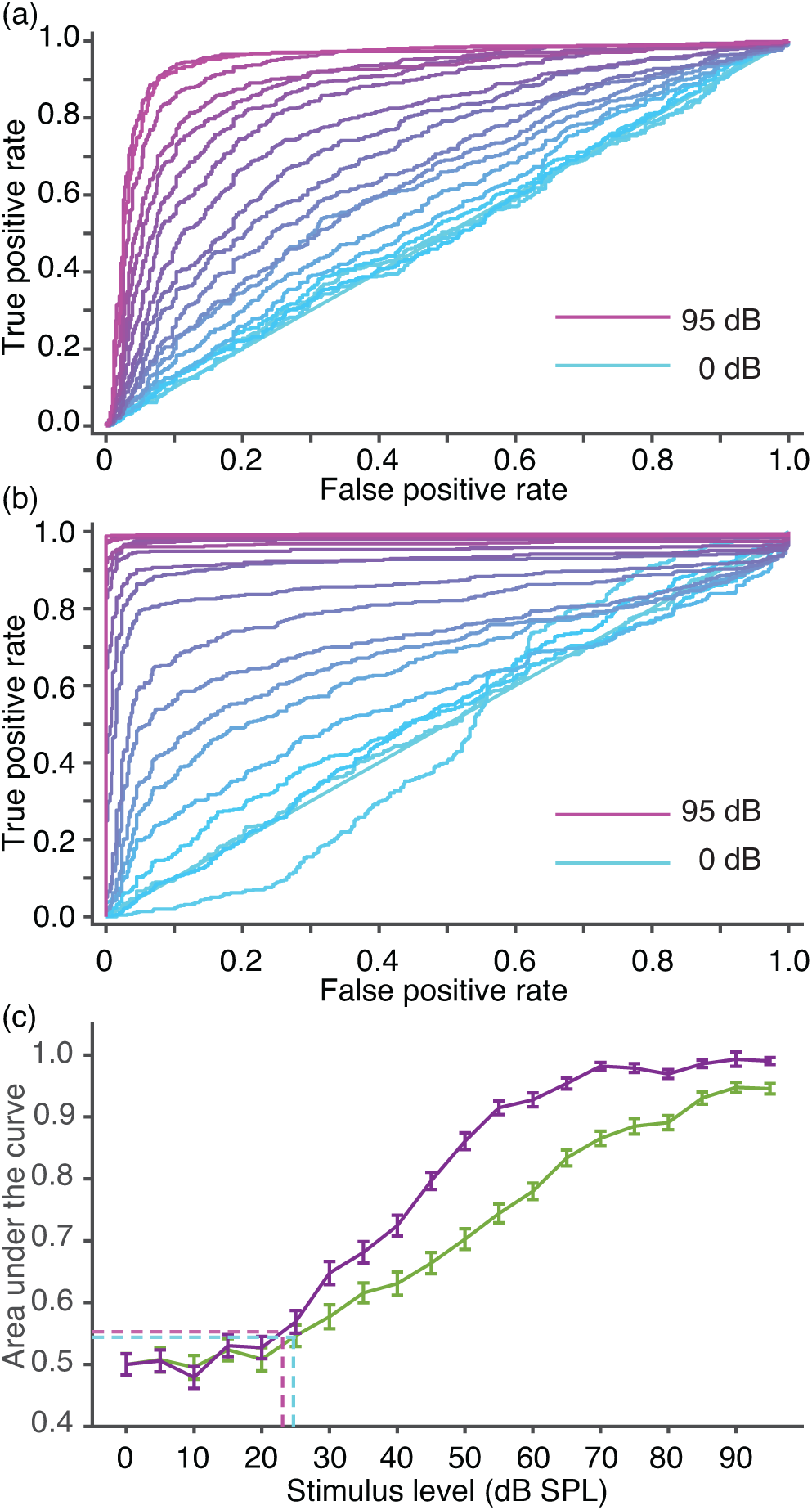
(a, b) ROC curves are shown in 5 dB increments for stimuli ranging from 0 dB SPL to 95 dB SPL. Error bars are omitted for clarity but are similar in size to those in Figs. 3 and 5. (a) ROC curves are shown for the voltage difference. (b) ROC curves are shown for the inner product. (d) The AUC is shown as a function of the stimulus level for the voltage difference (green) and the inner product (purple). The ABR threshold defined using the Mann-Whitney U test corresponds to an AUC larger than a lower bound set by Eqs. 5 and 13. For the voltage difference, an AUC of 0.54 yields an ABR threshold of 25 dB SPL (cyan dashed lines). When the AUC equals 0.55, the ABR threshold using the inner product is 23 dB SPL (magenta dashed line).

The ROC curves illustrate that perfect detection is almost possible at 95 dB SPL for the inner product but not for the voltage difference. Note that the inner-product ROC curves at 25 dB SPL and above drop below the line of no discrimination at high FPRs because their feature variance is larger than the no-stimulus feature variance (Fig 4)(Macmillan and Creelman, 2005). To compare the performance of the voltage difference with the inner product over all cutoff values, we employ a summary statistic.

The area under each ROC curve (AUC) given by

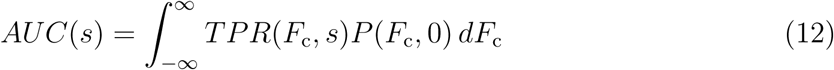

and equals the probability that a stimulus of size *s* evokes a feature value larger than feature values arising from no stimulus (Green and Swets, 1966; Fawcett, 2006). The AUC is independent of the number of stimulus presentations *N* and the feature cutoff. Larger AUCs correspond to better discrimination between the stimulus and no-stimulus conditions. At their respective PPTs, the voltage difference’s AUC is 0.89 ± 0.01 and the inner product’s is 0.73 ± 0.02. Although the values of the FPR and the TPR are the same for each feature at their PPTs, their AUCs differ (Welch’s t-test: *p <* 2 × 10*^−^*^14^). At and above 29 dB SPL, the inner product’s AUC exceeds that of the voltage difference (Welch’s t-test: *p* ≤ 0.05) (Fig. 6(d)). The features’ AUCs become indistinguishable as the level rises, however, because they converge to unity for high-intensity stimuli.

### ABD Threshold Defined by the Area Under the Curve

The PPT is based on the assumption that at threshold the auditory system should be able to set a cutoff ensuring a small FPR and a large TPR. Depending on how the auditory system is optimized, alternative definitions for an ABD threshold are possible. The AUC threshold can be defined as the sound pressure level at which the AUC achieves a particular value. For example, the stimulus level at which the AUC = 0.75 is 56 ± 2 dB SPL for the voltage difference and 42 ± 1 dB SPL for the inner product. A sufficiently large effect-size difference between the signal and noise distributions defines the ABD threshold.

### The ABR Threshold Defined by the Mann-Whitney U Test

The AUC can be thought of as the effect size of the Mann-Whitney U test as it is related to the Mann-Whitney U statistic *U* (*s*) according to

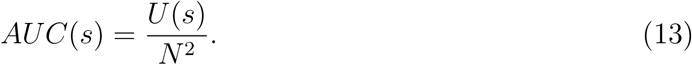

We use the Mann-Whitney U test to test the hypothesis that the feature distribution *P* (*F, s*) arising from a stimulus of size *s* is the same as the distribution in the absence of a stimulus *P* (*F,* 0) against the alternative that the distributions are different (Eq. 5). The inner-product threshold of 23 dB SPL is slightly lower than the voltage-difference threshold of 25 dB SPL (Fig. 6(d)). Because the U statistic depends on the number of stimulus presentations, the Mann-Whitney U threshold decreases as *N* increases. Although the Mann-Whitney U test does not compare average ABRs, it is an alternative to Welch’s t-test for defining the ABR threshold. Note that the ABR thresholds found using Welch’s t-test are similar to those found using the Mann-Whitney U test. The Mann-Whitney U threshold corresponds to a stimulus level at which a statistical difference can be found between a feature’s distribution and the no-stimulus distribution, whereas the AUC threshold is the level at which a detection effect size has been achieved.

### Signal-Detection Performance for a Population

To determine whether our observations hold more generally, we measured responses from four additional rats (Fig. 7). Across the population of five rats, the ABR thresholds of the voltage difference are similar to those of the inner product for both Welch’s t-test and the Mann-Whitney U test. FPRs are, however, large at the ABR thresholds found using Welch’s t-test for all rats. Over the population, the FPRs of the voltage difference (0.33 — 0.43) are larger than those of the inner product (0.23 — 0.37) (variance-reciprocal-weighted Welch’s t-test, *p <* 0.01, (Goldberg et al., 2005)). None of the feature distributions are normal (Q-Q plots indicate heavier tails than normal distributions, Anderson-Darling tests: *p <* 5 × 10*^−^*^4^, kurtosis = 4.6 — 54). PPTs are respectively 47 dB and 21 dB greater than ABR thresholds on average for the voltage difference and the inner product. Similar to the representative rat, the PPTs are 30 dB smaller for the inner product than they are for the voltage difference on average. The AUC threshold is more than 20 dB smaller for the inner product in comparison to the voltage difference. Finally, the inner product is better on average than the voltage difference at detecting the stimulus for sound pressure levels below 90 dB SPL and above 36 dB SPL according to the AUC (Fig. 8(a)).

**Figure 7:**
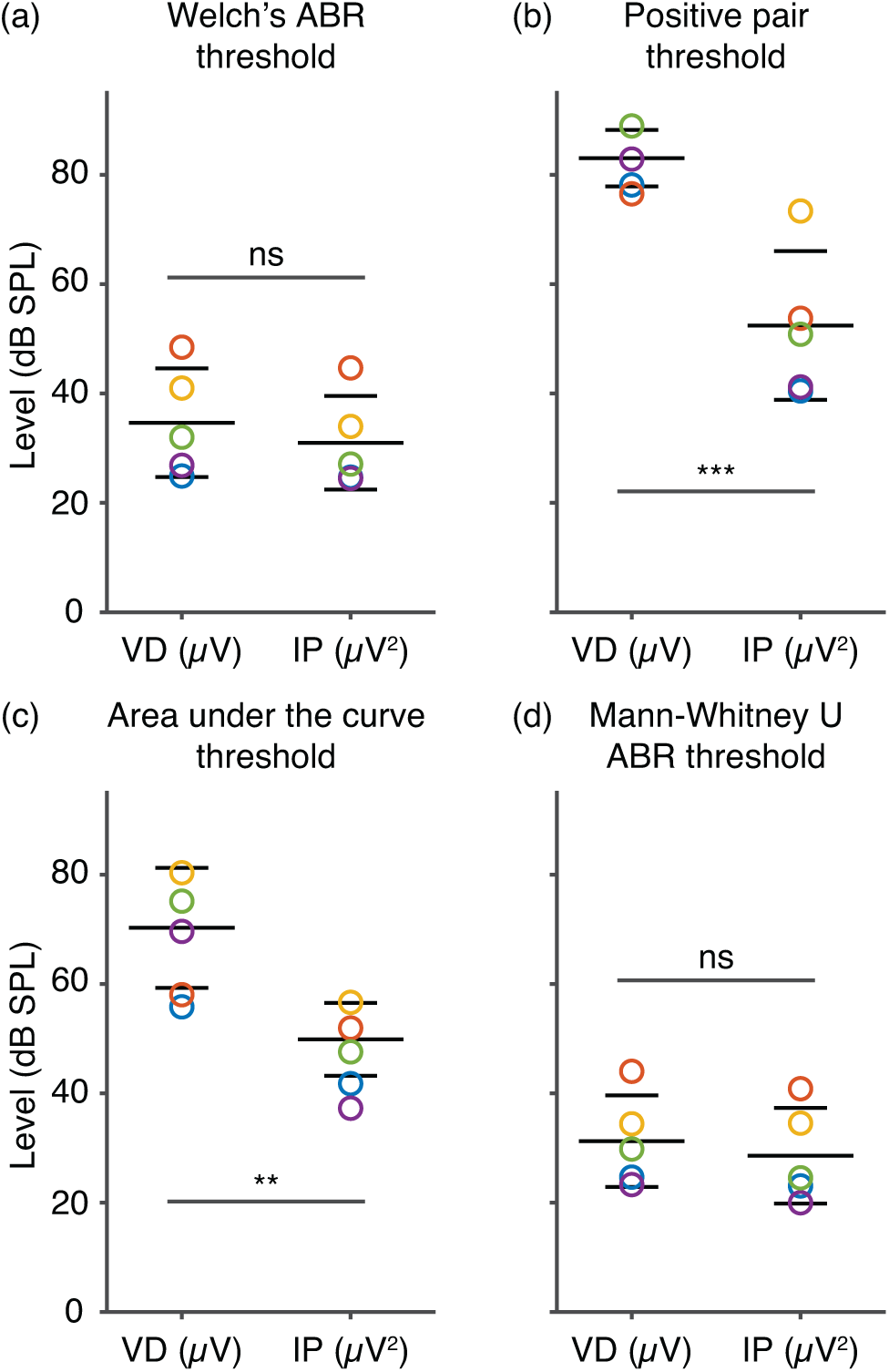
Measurements illustrating the ability of five rats to detect tone pulses are shown for the voltage-difference (VD) and the inner-product (IP) features. Each color indicates an individual rat’s results. The blue circles correspond to the rat discussed in Figs. 1-6. Bars indicate the weighted mean and one weighted standard deviation above and below the mean, in which the weights are reciprocals of the squared standard errors for each rat. Differences between the results for each feature are assessed using a weighted version of Welch’s t-test (ns – p-value *>* 0.05; * – p-value ≤ 0.05, ** – p-value ≤ 0.01, *** – p-value ≤ 0.005) (Goldberg et al., 2005). (a) The ABR thresholds using Welch’s t-test of the voltage difference are close to those of the inner product (p-value = 0.55). The inner-product blue circle is beneath the purple one. (b) The inner product’s PPT is 30 dB lower than that of the voltage difference on average. Because one rat’s TPR for the voltage difference failed to reach 0.5 for any stimulus level tested, this rat’s PPT could not be determined (The voltage-difference yellow circle is missing.). (c) The average level at which the AUC = 0.75 is smaller by 20 dB for the inner product in comparison to the voltage difference. (d) The ABR thresholds using the Mann-Whitney U test of the voltage difference are close to those of the inner product (p-value = 0.55).

**Figure 8:**
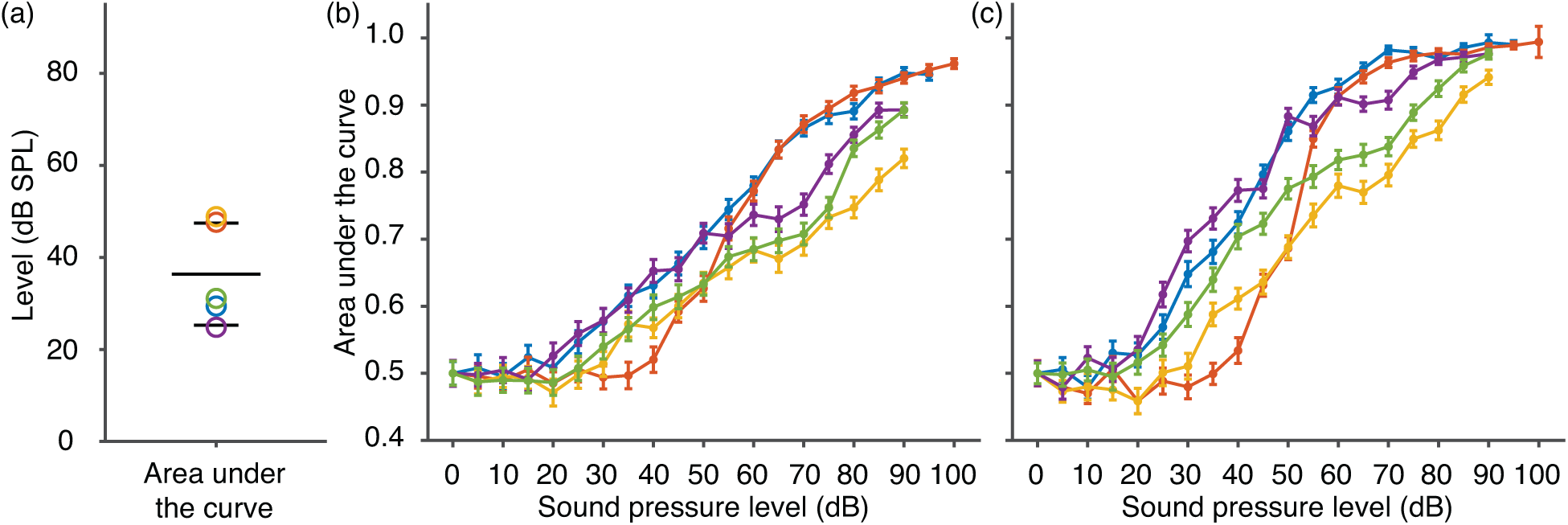
Supra-threshold ability of five rats to detect tone pulses are shown for the voltage-difference and the inner-product features. Each color indicates an individual rat’s results. The blue symbols correspond to the rat discussed in Figs. 1-6. (a) Up to at least 90 dB SPL for each rat, the levels at and above which the inner product’s AUC exceeds that of the voltage difference are shown (Welch’s t-test, *p* = 0.05). Bars indicate the weighted mean and one weighted standard deviation above and below the mean, in which the weights are reciprocals of the squared standard errors for each rat. (b) AUCs are shown as functions of the stimulus level for the voltage difference. (c) AUCs are shown as functions of the stimulus level for the inner product.

Supra-threshold signal-detection performance can be quantified by the AUC for all subjects (Fig. 8(b, c)). At each level, subjects can be ranked according to their hearing ability if we assign the same rank to rats lacking a statistically-significant difference. Because the AUC increases with stimulus level in a distinct manner for each rat, however, this ranking changes with level. A subject with relatively poor hearing at low levels may have better hearing than most of their peers at higher levels (red lines). At several stimulus levels, supra-threshold hearing deficits are indicated by the small AUCs of some subjects relative to their peers. Finally, some subjects with similar ABR thresholds exhibit very different supra-threshold hearing abilities. For example, the green and blue labeled inner product ABR thresholds are similar but the corresponding AUCs clearly differ between 45 and 80 dB SPL (Figs. 7(a,d) and 8(d)).

## Discussion

Here we show that trial-by-trial classification of ABRs is possible over all stimulus levels. In particular, we find ABD thresholds that are analogous to auditory thresholds found using behavioral methods. In each case, a determination is made about whether an individual stimulus presentation has been detected *before* the next stimulus is presented. In contrast, the ABR threshold is similar to defining threshold by asking a subject to decide once only *after* a set of stimuli was presented whether they heard the stimuli on average. This distinction is made all the more clear by considering the distribution of an ABR feature. The ABR threshold is defined by finding a statistically significant difference between the no-stimulus and the threshold distributions. In contrast, the ABD threshold is determined by a sufficiently large effect size based on the two distributions. Because effect sizes can only be reliably measured for statistically-significant differences, ABD thresholds are always larger than ABR thresholds. ABR thresholds based on finding statistically-significant differences between distributions, a temporal pattern in the average ABR, or signal-to-noise criteria decline as the number of stimulus presentations rises (Lütkenhöner and Seither-Preisler, 2008; Elberling and Don, 1984; Don et al., 1984), whereas the ABD threshold does not depend on the number of stimulus presentations.

There have been many efforts to relate ABR thresholds to behavioral thresholds (van der Drift et al., 1987; Stapells and Oates, 1997; Stapells, 2000; Gorga et al., 2006; Heffner et al., 2008; Borg, 1982; Longenecker et al., 2016; Walter et al., 2012; Ramos et al., 2013; Johnson et al., 2014). Although these empirical relationships are useful, their interpretation remains unclear, because behavioral thresholds answer a different question than that addressed by ABR thresholds, ABRs are limited to the brainstem, ABRs possess measurement noise, and the ABR is a poor representation of the neural signal. Determining how ABR trial-by-trial detection is related to behavior tests is far beyond the scope of this work. We can, however, make the following observations. First, trial-by-trial ABR classification only assesses the performance of the brainstem, whereas behavioral tests apply to the auditory system as a whole. Consequently, we should not expect trial-by-trial ABR classification to agree exactly with behavioral tests. Second, behavioral thresholds in rodents usually require stimuli with long durations (several seconds) or high intensities (over 80 dB SPL), which confounds comparison with ABD thresholds (Heffner et al., 2008; Borg, 1982; Longenecker et al., 2016; Koch and Schnitzler, 1997; B-laszczyk, 2003; Radziwon et al., 2009; Walter et al., 2012). Human behavioral thresholds decrease with increasing pure-tone stimulus durations, however, and plateau for sufficiently long durations (Zwislocki, 1960; Watson and Gengel, 1969; Moore, 2013; Pedersen and Elberling, 1972). Owing to the brief duration of the ABR stimulus, we expect the ABD threshold to be higher than pure-tone thresholds, for which the stimulus duration is 1 s or more (Gelfand, 2009).

The amplitude and latency of waves are effect sizes that could be used to define threshold or supra-threshold signal-detection performance. There is, however, considerable heterogeneity in amplitudes and latencies in the normal-hearing population (Lewis et al., 2015). Amplitude ratios and inter-peak latencies account somewhat for inter-subject variability, but waves are not apparent at low levels or in all subjects (Markand, 1994; Lütkenhöner and Seither-Preisler, 2008; Lewis et al., 2015). The trial-by-trial classification method described here accounts for inter-subject variability by comparing each subject’s response to their intrinsic noise background and provides signal-detection measures that can be compared across subjects. We can use these measures to rank the hearing ability of normal hearing subjects at supra-threshold levels. Trial-by-trial classification can be done for response features at all stimulus levels, allowing us to circumvent wave-identification difficulties. We find that supra-threshold hearing performance improves with rising stimulus levels in a subject-dependent manner. At each stimulus level, small AUC values for some subjects relative to their peers indicates these rats possess supra-threshold hearing deficits, possibly arising from a loss of high-threshold auditory nerve fibers, despite having normal ABR thresholds (Liberman, 1978; Kujawa and Liberman, 2009; Schaette and McAlpine, 2011).

A subject’s responses in a behavioral test depend not only on their hearing ability but also on their attentional state, which might vary during the test. Because we measure a feature’s distribution directly, we avoid attentional bias in estimating decision distributions (Macmillan and Creelman, 2005). In signal-detection theory, an attentional level is defined by a feature’s cutoff and determines the FPR and TPR. To address attentional changes, we define threshold in terms of a small FPR and a large TPR, the positive pair (FPR, TPR) threshold, or quantify hearing ability across attentional states using the AUC. We find, the same positive pair (FPR, TPR) corresponds to very different AUC values for the voltage difference in comparison to the inner product. These differences arise from the two features having distributions that differ qualitatively. Whether it is appropriate to define the ABD threshold in terms of the pair (FPR, TPR), the AUC, or other signal-detection measures depends on what characteristics the auditory system has optimized (Powers, 2011).

The voltage difference is defined using the average ABR and is equivalent to the peak-to-trough difference of Jewett waves at some stimulus levels (Jewett et al., 1970; Jewett, 1970; Jewett and Williston, 1971; Buchwald and Huang, 1975). Although the voltage difference measures the largest peak-to-trough difference, its physiological interpretation is less clear than that of a Jewett wave. To determine the ABD threshold for a Jewett wave, however, the wave must be defined for stimulus levels at which the waveform is unclear (Fig. 1). Nonetheless, studying ABD thresholds for features that overlap with Jewett waves may be fruitful.

The inner product is similar to crosscorrelation methods used to analyze ABRs. Cross-correlations between two average ABRs (Weber and Fletcher, 1980; Arnold, 1985; Mason et al., 1977; Mason, 1984; Picton et al., 1983; Valdés-Sosa et al., 1987; Mijares et al., 2013; Valderrama et al., 2014; Özdamar et al., 1994; Davey et al., 2007), the average ABR and a template based on the average ABR at certain stimulus levels (Salomon et al., 1973; Elberling, 1979; Cone-Wesson et al., 1997), and the average ABR and a template based on a set of normal ABRs across subjects (Neely and Pepe, 1997; Elberling, 1979; Valderrama et al., 2014), as well as the average correlation between pairs of individual ABRs (Coppola et al., 1978) have been employed. The average inner product has been introduced previously, but its use is not widespread (Coppola et al., 1978). We find the inner-product distribution to be particularly useful for summarizing the entire ABR epoch as it quantifies the similarity between and the magnitude of individual ABRs.

In the clinic, a single point of the average ABR, the amplitude of wave V, is typically used to assess hearing functionality (Burkard et al., 2007; Hall, 2007; Johnson et al., 2014). By comparing two features, the voltage difference and the inner product, we illustrate how using the entire ABR response time, yields improved signal-detection performance. If we had employed only the average ABR, we would have concluded that the two features were equivalent for determining the threshold of hearing owing to their ABR thresholds being similar. We find, however, that the inner product’s PPT is lower than that of the voltage difference by 30 dB and that the supra-threshold performance of the inner product surpasses that of the voltage difference for stimulus levels below 90 dB SPL. Because the inner product is based on all the points in an epoch whereas the voltage difference is defined by only two points, it is not surprising that the inner product outperforms the voltage difference. The effects of the background noise are somewhat ameliorated by taking all of the points in an epoch into account.

In psychophysical studies of threshold, intrinsic noise limits a subject’s sound-detection ability (Green, 1960; Green and Swets, 1966; Macmillan and Creelman, 2005). Because some sources of intrinsic ABR noise differ from those affecting the firing of neurons in the auditory pathway and ABRs possess measurement noise, ABR noise differs from behavioral noise (Elberling and Wahlgreen, 1985; Don and Elberling, 1994; Chesnaye, 2019). Thus the signal-detection task in a behavioral test cannot be identical to trial-by-trial ABR classification. Nonetheless, ABR classification performance differs between subjects and improves with increasing stimulus levels, implying this approach quantifies the signal-detection ability of the brainstem.

Because signal-detection is inherently stochastic, trial-by-trial classification of signals is frequently used to evaluate artificial and natural systems (Pernet et al., 2011; Powers, 2011; Trenado et al., 2019; Wilmshurst, 1990; Ye et al., 2018). The classification methodology we employ does not require training data and provides objective and interpretable measures of signal-detection performance that can be compared across systems. We show this approach is feasible in rodents, yields thresholds different to traditional ABR analysis techniques, and provides information about a subject’s supra-threshold hearing ability. Trial-by-trial ABR classification holds promise as a new method to detect hearing impairment as a function of stimulus level in animal models and in humans.

## Acknowledgements

We would like to thank Anthony J Ricci for providing resources and supporting Noor-E-Seher Ali through grant number 5P01AG051443 from the National Institute on Aging, a division of the National Institutes of Health. George Liu would like to thank Stephen Bates for discussions. We are grateful for funding from the Stanford Initiative to Cure Hearing Loss supported by a generous gift from the Bill and Susan Oberndorf Foundation and The Shari and Kenneth Eberts SICHL Research Fund. We are also thankful for support from the Anna and Nathan Flax Foundation.

